# Aerobic exercise promotes PDAC vascular normalization through S1PR1 signaling in tumor endothelial cells

**DOI:** 10.1101/2024.02.14.580332

**Authors:** Riccardo Ballarò, Sumedha Pareek, Jonghae Lee, Hetal Patel, Priya Tirumala, Guanshu Liu, Karma Hayek, Hannah Savage, Bella Guerrouahen, An Ngo-Huang, Nathan Parker, Matthew HG Katz, Maria Petzel, Florencia McAllister, Robin Wright, Lin Tan, Philip L Lorenzi, Keri Schadler

## Abstract

Pancreatic Ductal Adenocarcinoma (PDAC) is characterized by high resistance to anti-cancer therapies. This resistance is caused in part by hypo-vascularization and dysfunctional vessels which inefficiently deliver chemotherapy. Here, we define the mechanism of aerobic exercise-induced tumor vascular remodeling and improved chemotherapy delivery and efficacy. We found that aerobic exercise was able to improve tumor vascular function and increase the number of lymphatic vessels, and these effects were associated with increased gemcitabine delivery to and efficacy against PDAC in mice. Further, exercise increased sphingosine-1-phosphate (S1P) in plasma and the activation of sphingosine-1-phosphate receptor 1 (S1PR1) in tumor endothelial cells. S1PR1 endothelial cell-specific deletion blunted the exercise-induced improvements in tumor vascular function, gemcitabine efficacy and drug concentration. Preclinical findings were validated in a patient cohort in which we found that exercise during neoadjuvant chemotherapy remodeled PDAC vasculature and improved tumor vascular function. These findings provide direct evidence that exercise increases chemotherapy delivery and efficacy by improving vascular function, defining S1PR1 as a necessary mediator of exercise-induced vascular remodeling.

## Introduction

Pancreatic Ductal Adenocarcinoma (PDAC) is the third leading cause of cancer-related death in the Unites States, with an overall 5-year survival of 12%^1^. Although the recent treatment advances for PDAC have improved the outcome of resectable localized tumors, approximately 80% of patients with PDAC are not candidates for curative surgery^2^. Various combinations of systemic chemotherapy with or without radiation have been used over the past several decades with little progress toward better outcomes^3^. These poor results reflect the aggressive biology of PDAC cells, along with the complex tumor microenvironment which promotes cancer proliferation, metastasis and therapeutic resistance^4^. The PDAC microenvironment is composed of poorly formed and dysfunctional vessels, cancer-associated fibroblasts, dense extracellular matrix, and largely suppressive immune cell populations^5^. Although dysfunctional vasculature is a common feature of most solid tumors, PDAC vasculature is particularly insufficient for drug delivery to the tumor because most PDAC tumors have low abundance of blood vessels in addition to the dysfunction of existing vessels^6^. The vascular dysfunction is compounded by the dense stroma, which increases intratumoral solid stress, causing vessel collapse^7^. These vascular alterations, together with insufficient lymphatic drainage, result in high interstitial fluid pressure, ultimately preventing drugs from accessing the tumor^8–10^. Promoting the formation of functional vessels may be an effective strategy to increase chemotherapy efficacy and decrease cancer dissemination.

We previously demonstrated that aerobic exercise positively impacts tumor vasculature in mice^11–14^. Exercise remodeled melanoma vasculature, increasing microvessel density, the number of long vessels and vessels with open lumens^11,15^. Similarly, aerobic exercise improved Ewing sarcoma-bearing mouse vasculature and chemotherapy efficacy and these effects were associated with increased S1PR1 expression in tumor endothelial cells^14^. The exercise-induced remodeling of tumor vasculature was also observed in a subcutaneous PDAC patient-derived xenograft and syngeneic PDAC cell line in mice^11,12^. However, the subcutaneous location of these tumor models is unlikely to reflect the correct tumor microenvironment, making them inferior models for defining the relationship between exercise and PDAC tumor vasculature. Here, we evaluate the effects of aerobic exercise on orthotopic PDAC vasculature and response to therapy. Further, we utilize an orthotopic PDAC model to define the molecular regulation of the tumor vascular response to exercise.

Vascular maturation and function are regulated by a family of G-coupled receptors, the S1P receptors (S1PRs), which are activated by the ligand sphingosine-1 phosphate (S1P)^16^. S1PR1, S1PR2 and S1PR3 are expressed by endothelial cells and together regulate vascular development, tone and permeability^16^. S1PR1 signaling in response to S1P suppresses VEGF-dependent vascular sprouting, induces adherens junction assembly, and enhances integrin activation as a fundamental mechanism for vascular stabilization and efficient perfusion^17^. In addition, S1PR1 regulates the interaction of perivascular and endothelial cells in the microvasculature to generate mature, functional vessels^18^. Deletion of the *s1pr1* gene is embryonically lethal due to defective neovascularization^19,20^. Post-natal endothelial cell-specific deletion of *s1pr1* impairs vascular architecture, permeability and barrier function^21^. S1PR1 has also been shown to be an important regulator of tumor vessel function^22,23^.

We demonstrate that exercise improves tumor blood and lymphatic vessels, and that the remodeling of PDAC vasculature by exercise is associated with increased plasma S1P and S1PR1 activation in tumor vascular endothelial cells. Exercise also reduced peritoneal spreading of PDAC. Our findings show that S1PR1 expression is necessary for the tumor vascular improvements and for the increased chemotherapy delivery and efficacy induced by exercise in PDAC-bearing mice, establishing a causal effect between improved tumor vasculature and increased chemotherapy response in tumors upon exercise. Further, we validate the relationship between exercise and vasculature in a human PDAC dataset, showing that increased physical activity in PDAC patients remodels tumor vasculature and reduces tumor vascular leakage.

## Methods

### Cell culture

PDAC KPC-4662 cells were derived from *Kras*^LSL-G12D/+^; *Trp53*^LSL-R172H/+^; *Pdx1-Cre* mice and provided by Dr. Robert Vonderheide (University of Pennsylvania School of Medicine). Hy 15549 cells were derived from *Kras*^LSL-G12D/+^; Trp53*^lox/+^; Ptf1a*-*Cre* mice and provided by Dr. Haoqiang Ying (The University of Texas MD Anderson Cancer Center). Cells were cultured in DMEM (Gibco 11965092; KPC-4662) or RPMI (Corning 10-040-CV; Hy15549) containing 10% FBS (Sigma-Aldrich, F2442), 100 U/mL penicillin, and 100 mg/mL streptomycin (Gibco, 15140122). Cells were passaged by dissociation with Accutase (Sigma-Aldrich, A6964) and maintained at 37°C, 95% relative humidity, and 5% CO_2_. Cells were routinely confirmed as Mycoplasma negative by qRT-PCR with primers forward 5’-GGGAGCAAACAGGATTAGATACCCT-3’ and reverse 5’-TGCACCATCTGTCACTCTGTTAACCTC-3’, or by analysis with MycoAlert Mycolplasma Detection Kit (Lonza, LT07-418).

### Mouse strains

The Institutional Animal Care and Use Committee at The University of Texas MD Anderson Cancer Center approved all animal studies. Mice were housed in cohorts of three to five animals per cage, in individually ventilated cages under pathogen-free conditions. Animals had free access to food and water and were kept on a 12-hour light/12-hour dark cycle. C57BL/6J wild-type mice were purchased from The MD Anderson Experimental Radiation Oncology Mouse Facility or Jackson Laboratory. S1PR1 GFP signaling reporter mice were previously described^24^. Briefly, B6N.129S6(FVB)-*S1pr1*^tm3.1(tTA,-Arrb2)Rlp/J^ mice (Jackson Laboratory, stock #026275) were crossed with pTRE-H2BGFP mice (Jackson Laboratory, stock #005104) for the generation of a mouse carrying a S1PR1 knock-in with tetracycline-regulated transactivator (tTA)-tobacco etch virus (TEV) protease recognition sequence (tevs) fusion protein and an *Arrb2* (murine β-arrestin)-TEV protease fusion protein, and the transgene expressing the human histone 1 (H2BJ) protein and Green Fluorescent Protein (GFP) fusion protein under the control of a tetracycline-responsive promoter element (TRE; tetO). pTRE-H2BGFP were purchased with a BALB/c background and were backcrossed 7 times with C57BL/6J mice before being used for experiments. Mice expressing one allele of both transgenes were considered S1PR1 GFP reporter mice. Littermates expressing only the H2BJ-GFP allele without the S1PR1 knock-in were used as controls^24^. Endothelial cell-specific *S1pr1* knockout mice (*S1pr1*^LoxP/LoxP^ *Cdh5*-*Cre*-ER^T2^; S1PR1 EKO) were previously described^20^. Briefly, mice carrying *loxP* sites flanking *S1pr1* gene (*S1pr1*^loxP/loxP^; S1PR1 WT; Jackson Laboratory, stock # 019141) were crossed with mice expressing Cre recombinase (*Cre*-ER^T2^) under the vascular endothelial cadherin (*Cdh5*) promoter (*Cdh5*-*Cre*ER^T2^; Taconic # 13073). Gene deletion by the Cre recombinase was achieved with tamoxifen (Sigma-Aldrich, T5648; 100 mg/kg per day) treatment by oral gavage at 6-9 weeks of age for five consecutive days, and mice were allowed to recover for 12 days before being used for experiments. Mouse genotyping was performed using ear punches. The genotype and phenotype of these mice was verified by analyzing the DNA recombination in lungs and by immunofluorescence in tumor tissue, respectively (**Supplementary** Fig. 1a, b).

### Tumor cell injection, experimental protocol and exercise regimen in mice

KPC-4662 (1.25 x 10^5^ suspended in cell medium) or Hy 15546 cells (1 x 10^4^ suspended in Hank’s Buffered Saline Solution) were injected into the mouse pancreas in 20µl volume, during aseptic surgery. Briefly, an incision was made in the left flank and tumor cells were injected using a 30-gauge needle into the tail of the pancreas. The peritoneum was sutured with 4-0 Vicryl and the skin was closed with wound clips. To calculate the tumor volume, two tangential ultrasound images along the longitudinal and transverse planes were used. Tumor volume was calculated by the formula 4/3πABC, where A, B and C are the measured diameters of each tumor using NIH open-source image software ImageJ (http://rsbweb.nih.gov/ij/; **Supplementary** Figure 1c). Tumor volumes were then measured once per week by the same method.

### Experimental protocol 1: exercise 8 and 16 meters per minute (m/min)

At day 8 after KPC-4662 or Hy 15549 injection, tumor volume was measured by ultrasound using Vevo 3100 with a 40-MHz linear signal transducer and 550D probe (Fujifilm, VisualSonics), and mice were divided by tumor size to the different experimental groups: sedentary (Sed; n= 9 KCP-4662; n=10 Hy 15549), exercise 8 m/min (Ex 8; n=9 KPC-4662; n=10 Hy 15549) and exercise 16 m/min (Ex 16; n=9 KPC-4662; n=10 Hy 15549). Vascular function (vessel perfusion and leakage) and hypoxia in KPC-4662 mice were assessed in different experiments using the same conditions except for the experimental groups, which included only Sed (n=11-10) and Ex 16 (n=10). Mice underwent the acclimatization protocol on treadmills (Exer 3/6 Treadmill; Columbus Instruments) at day 9 and 10 after tumor injection. For the 8 m/min exercise protocol, acclimatization consisted of exercise at 3 m/min for 20 minutes on day 9, and at 6 m/min for 40 minutes on day 10 (**Supplementary** Figure 1d). For the 16 m/min exercise protocol, on day 9 mice ran at 8 m/min for 20 minutes and on day 10 at 12 m/min for 40 minutes. Mice started the full exercise regimen after tumors reached ∼ 30 mm^2^ (day 12; 8 m/min or 16 m/min for 45 minutes, 5 days a week for around 3 weeks; **Supplementary** Figure 1d). All mice were humanely euthanized at day 29-31 by cervical dislocation under isoflurane anesthesia 48 hours after the final exercise session.

### Experimental protocol 2: exercise 16 m/min in S1PR1 GFP reporter mice

KPC-4662 cells were injected as described above into S1PR1 GFP reporter mice (B6N.129S6(FVB)-*S1pr1*^tm3.1(tTA,-Arrb2)Rlp/J^ x pTRE-H2BGFP mice). At day 8 after KPC-4662 cell injection, were divided into two groups: Sed (n=7) or Ex (exercise 16 m/min; n=7). Mice underwent the acclimatization protocol and full exercise regimen as described above. pTRE-H2BGFP mice (n=2) were also added to the experiment as a negative control for GFP fluorescence. Only mice that survived until the end of the experimental protocol were used for flow cytometry analysis of S1PR1 activation (Sed, n=6; Ex, n=5). All mice were humanely euthanized as described above.

### Experimental protocol 3: exercise 16 m/min + gemcitabine in S1PR1 WT mice

At day 8 after KPC-4662 cell injection, *S1pr1*^loxP/loxP^ (S1PR1 WT) mice were randomized in Sed (n=11), Ex (exercise 16 m/min; n=11), Gem (gemcitabine; n=11) and Ex+Gem (exercise+gemcitabine; n=11). Mice underwent the acclimatization protocol and full exercise regimen as described above. Mice were treated with intravenous (i.v.) injections of 25 mg/kg gemcitabine (Sagent Pharmaceuticals, 25021-235-50) after the exercise session, twice per week, for 3 weeks). All mice were humanely euthanized as described above.

### Experimental protocol 4: exercise 16 m/min + gemcitabine in S1PR1 WT and S1PR1 EKO mice

Both S1PR1 WT (*S1pr1*^loxP/loxP^) and S1PR1 EKO (*S1pr1*^LoxP/LoxP^ *Cdh5*-*Cre*-ER^T2^) mice were treated with tamoxifen as described above. At day 8 after KPC-4662 cell injection, S1PR1 EKO mice were randomized in Sed (n=10), Ex (exercise 16 m/min; n=10), Gem (gemcitabine; n=11) and Ex+Gem (exercise+gemcitabine; n=11). S1PR1 WT mice (n=11) were also added to the experiment as a control and for the evaluation of basal differences between S1PR1 and EKO groups. Mice underwent the acclimatization protocol, full exercise regimen, chemotherapy treatment and terminal euthanasia as described above.

### Peritoneal carcinomatosis (PCI) analysis

The PCI analysis was performed using a scoring system adapted from Bastiaenen *et al.*^25^. At the time of necropsy, mice were evaluated for the presence of macroscopic metastases in different areas and organs of the peritoneum (liver, diaphragm, spleen, intestine, peritoneum wall). Per organ or region, a score of 0-3 was determined based on the number and size of nodules. No macroscopic metastasis was scored as 0; tumor ≤ 2mm was scored as 1; tumors 2-5mm or 2 - 5 tumors was scored as 2; tumor ≥ 5mm or > 10 tumors was scored as 3. The PCI of each animal was calculated by adding scores of all the different regions within the animal together.

### Immunofluorescence Staining

For frozen sections of mouse tumors, tissues were fixed in 4% PFA for 1 hour at 4°C. Tissues were washed three times in PBS, incubated for 48 hours with rocking at 4°C in 30% sucrose and embedded in OCT compound. Frozen sections were cut at 10 µm thickness. Tissue sections were fixed with cold acetone for 8 minutes at −20°C, then washed three times in PBS before blocking with 4% fish gel blocking solution (CWFS 900.0033, Aurion) in PBS for 1 hour at room temperature. Primary antibodies were diluted in blocking solution and incubated at 4°C overnight. Slides were washed three times with PBS after the overnight incubation. Secondary conjugated antibodies were diluted in blocking solution and incubated for 1 hour at room temperature. Slides were mounted with Fluoro-Gel II with DAPI solution to stain the nuclei (Electron Microscopy Science, 17985-50). The following antibodies and dilutions were used: rat anti-mouse CD31 1:50 (BD Biosciences, 550274), goat anti-mouse CD31 1:50 (R&D Systems, AF3628), rat anti-mouse LYVE-1 1:50 (Santa Cruz Biotechnology, sc-65647), rabbit anti-mouse Albumin 1:100 (Proteintech, 16475-1-AP), rabbit anti-mouse/human αSMA 1:100 (Abcam, ab5694), rabbit anti-mouse NG2 1:200 (EMD Millipore, AB5320), rabbit anti-mouse Desmin 1:200 (Abcam, ab15200), rat anti-mouse S1PR1 1:50 (R&D Systems, MAB7089). The following secondary antibodies and dilutions were used: goat anti-rat Alexa Fluor 488 1:1000 (Thermo Fisher Scientific, A-11006), goat anti-rat Alexa Fluor 594 1:1000 (Thermo Fisher Scientific, A-11007), donkey anti-rabbit Alexa Fluor 647 1:1000 (Thermo Fisher Scientific, A-31573), goat anti-rabbit Texas Red 1:1000 (Thermo Fisher Scientific, T-2767), donkey anti-goat 647 1:1000 (Thermo Fisher Scientific, A21447).

For human tissue, paraffin-embedded slides were deparaffinized using xylenes and ethanol. Antigen retrieval was performed in 10mM Tris Base, 1mM EDTA solution for 20 minutes. Slides were washed three times in PBS before blocking with 4% fish gel blocking solution (CWFS 900.0033, Aurion) in PBS for 30 minutes at room temperature. Primary antibodies were diluted in blocking solution and incubated at 4°C overnight. Slides were washed three times with PBS after the overnight incubation. The following primary antibodies and dilutions were used: rabbit anti-human CD31 1:50 (Abcam, ab28364) and goat anti-human Albumin 1:50 (Abcam, ab19194). Secondary conjugated antibodies were diluted in blocking solution and incubated for 1 hour and 30 minutes each at room temperature. Secondary incubation for the two different secondaries used was performed in two separate steps of 1 hour and 30 minutes each, spaced out by three PBS washes. The following secondary antibodies and dilutions were used: donkey anti-goat 647 1:500 (Thermo Fisher Scientific, A21447) and goat anti-rabbit 594 1:1000 (Thermo Fisher Scientific, A-11012). Slides were mounted with Fluoro-Gel II with DAPI solution to stain the nuclei (Electron Microscopy Science, 17985-50).

### Tumor hypoxia and vessel perfusion

To evaluate the number of perfused vessels, mice were injected via tail vein with 100 µL Lycopersicon esculentum (Tomato) Lectin-LEA DyLight (1 mg/mL; Invitrogen, L32471) and euthanized after ten minutes. Tumors were excised and processed for immunofluorescent staining as described above. For tumor hypoxia analysis, mice were injected via tail vein with 100µL Hypoxyprobe-1 (pimonidazole HCl; 18 mg/ml; Hypoxyprobe Inc., HP6-200Kit) and euthanized after one hour. Tumors were processed for immunofluorescence staining and incubated with FITC conjugated to mouse IgG1 monoclonal antibody (FITC-Mab1; Hypoxyprobe Inc, 1:75) diluted in 4% fish gel blocking solution overnight at 4C. Whole tumor tile scan images were captured using Leica LAS X software. The hypoxia area of tumor sections was calculated by normalizing hypoxyprobe-1^+^ area to total number of nuclei. Area of total lectin^+^ signal was normalized by total CD31^+^ area. Sample numerosity for these analyses may vary from the original mouse number because for very ill mice, i.v. injection of the hypoxyprobe-1 or lectin was deemed to be ineffective and therefore hypoxia and vessel perfusion were not analyzed in those mice.

### Tumor vascular image analysis

Five to nine images were randomly captured at 10x magnification across each tumor and stained as described above. Quantifications were performed on each image and all image quantifications corresponding to a sample were averaged to report one value per tumor on graphs. Images were captured using a Leica fluorescence microscope (DMi8; Leica Microsystems Inc.) with LAS X software. Image analyses were performed using ImageJ. Tumor vessel structural analysis was performed as described previously^11^. Vessel parameters included the total number of CD31^+^ vessels, the percentage of visible open lumens over total vessels, the percentage of vessels with a length >100 µm (elongated vessels), and the number of 100 µm^2^ regions per image occupied by CD31^+^ vessels (microvessel density). Specifically, the microvessel density was calculated by dividing the image area into equal squares (100 µm^2^) creating a grid. The number of squares in the grid occupied by CD31^+^ vessels was counted, and total number of squares are reported as microvessel density. For evaluating the pericytes coverage of blood vessels, the pericyte markers αSMA, desmin or NG2 were costained with CD31. Pericytes coverage was quantified as the colocalization between pericyte markers and CD31 fluorescence area.

### Flow cytometry

KPC-4662 tumors from sedentary and exercised mice were harvested and immediately digested in HBSS with DNase I (100 µg/mL; Roche, 04716728001) and collagenase type II (10mg/ml; Worthington, LS004176) for 45 minutes at 37°C. Reb blood cells were lysed using Ammonium-Chloride-Potassium (ACK) lysing buffer (Thermofisher Scientific, A1049201) for 5 minutes on ice. Single-cell suspensions were prepared using a 100 µm nylon mesh. Cells were blocked for 15 minutes at room temperature using purified rat anti-mouse CD16/CD32 (BD Biosciences, 553141). Cells were stained with fluorochrome-conjugated antibodies for 15 minutes at room temperature, followed by dead cell staining with aqua fluorescent reactive dye (Invitrogen, 2456941) for 20 minutes and then fixed with stabilizing fixative (BD Biosciences, 338036). Data were acquired using an LSR Fortessa flow cytometer (BD Biosciences) and analyzed with FlowJo software version 10. Fluorochrome-conjugated antibodies used for flow cytometry were CD45 Pacific Blue (Biolegend, 103126) and CD31 PE-Cy7 (Invitrogen, 25-0311-82).

### Analysis of S1P by Triple Quadruple LC-MS

Blood from sedentary or exercised mice was collected by retro orbital bleeding in EDTA collection tubes 2 hours after the last bout of exercise. Blood was centrifuged twice at 2000 g for 10 minutes to pellet the cellular component and the supernatant was collected in a new tube after each centrifugation.

Analysis was performed by the Metabolomics Core Facility at MD Anderson Cancer Center. To determine the relative abundance of D-sphingosine (DSP) and S1P in mouse plasma samples, extracts were prepared and analyzed by TSQ Quantiva triple quadruple mass spectrometer coupled with a Dionex UltiMate 3000 HPLC system (Thermo Fisher Scientific). Metabolites were extracted using ice cold methanol with S1P-d7 as stable isotope internal standards. The lysates were vortexed, centrifuged at 17,000 g for 5 minutes at 4°C, and supernatants were transferred to clean autosampler injection vials and 10 μL was injected for DSP/S1P analysis by liquid chromatography (LC)-MS. The mobile phase A(MPA) is 0.1% Formic Acid in water. Mobile phase B(MPB) is 0.1% Formic Acid in acetonitrile.

Separations of S1P were achieved on an InertSustain Bio C18 HP, 3 µm, 100 x 2.1 mm column (GL Sciences). The flow rate was 200 µL/min at 35 °C, and the gradient elution program was: initial 40% MPB, increased to 60% MPB at 2 min, held at 60% MPB for 5 min, returned to initial conditions and equilibrated for 4 min. The total run time was 11 min. The mass spectrometer was operated in the MRM positive ion electrospray mode with the following transitions. S1P: m/z 380.3→264.2, DSP: m/z 300.3→252.3, S1P-d7 (IS): m/z 387.3 → 271.2, DSP-d7: 307.3 → 259.3. Raw data files were imported to Trace Finder software (Thermo Fisher Scientific) for final analysis. The relative abundance of S1P was normalized by internal standards.

### Analysis of gemcitabine by LC-MS

KPC-4662-bearing mice treated with gemcitabine (Gem) or with exercise plus gemcitabine (Ex+Gem) received a final i.v. dose of 25 mg/kg gemcitabine 1 hr prior to euthanization. Tumors were then collected and frozen in liquid nitrogen for further analysis. To determine the relative abundance of Gemcitabine (dFdC), extracts were prepared and analyzed by ultra-high-resolution mass spectrometry (HRMS). Approximately 20-30 mgs of tissue were pulverized on liquid nitrogen, then homogenized with Precellys Tissue Homogenizer (Bertin Corp.). Metabolites were extracted using 1 mL ice-cold 80/20 (v/v) acetonitrile/water. Extracts were centrifuged at 17,000 g for 5 min at 4°C, and supernatants were transferred to clean tubes, followed by evaporation to dryness under nitrogen. Dried extracts were reconstituted in 50/50 (v/v) acetonitrile/water, and 10 μL was injected for analysis by liquid chromatography (LC)-MS. LC mobile phase A (MPA) was 95/5 (v/v) water/acetonitrile containing 20mM ammonium acetate and 20mM ammonium hydroxide (pH∼9), and mobile phase B (MPB) was acetonitrile. Vanquish LC system (Thermo Fisher Scientific) included a Xbridge BEH Amide column (3.5 µm particle size, 100 x 4.6 mm) with column compartment kept at 35°C. The autosampler tray was chilled to 4°C. The mobile phase flow rate was 350 µL/min, and the gradient elution program was: 0-1 min, 85% MPB; 1-16 min, 85-5% MPB; 16-20 min, held at 5% MPB; 20-21 min, 5-85% MPB; 21-25 min, re-equilibrate column at 85%MPB. The total run time was 25 min. Data were acquired using an Orbitrap Exploris 240 Mass Spectrometer (Thermo Fisher Scientific) under ESI negative ionization mode at a resolution of 120,000 with Select Ion Monitoring (SIM) mode. Raw data files were imported to Trace Finder 5.1 software (Thermo Fisher Scientific) for final analysis. The relative abundance of each metabolite was normalized by tissue weight. Sample numerosity for this analysis may vary from the original mouse number due to the variable achievement of the final i.v. injections in mice that were very ill.

### DNA isolation and amplification for S1PR1 EC-KO characterization

The lungs of KPC-4662-bearing mice were harvested and immediately frozen in liquid nitrogen. DNA was extracted with DNAzol reagent (Invitrogen, 10503-027) following manufacture’s protocol. Briefly, lungs were homogenized in DNAzol reagent, centrifuged to remove insoluble tissue fragments and RNA, and DNA was precipitated and washed using ethanol 100% and 75%, respectively. DNA was then pelleted and resuspended in 8 mM NaOH, 1M HEPES, pH 7.2, at a concentration of 0.2-0.3 µg/µl. To detect the wild type (S1PR1 WT), knock-out (S1PR1 KO), and conditional alleles (S1PR1 loxP) by PCR the following primers were used: P1, 5’ GAGCGGAGGAAGTTAAAAGTG; P2, 5’ CCTCCTAAGAGATTGCAGCAA. P1 and P2 amplify an approximately 250-bp fragment for the S1PR1 loxP allele and a 200-bp fragment for the S1PR1 WT and S1PR1 KO alleles. To detect the recombined S1P1 allele, P1 and P3 (5’ GATCCTAAGGCAATGTCCTAGAATGGGACA) were used. P1 and P3 amplify a 200-bp fragment. When primers P1, P2, and P3 were used in the same PCR reaction, the PCR products were digested with SacI (Thermo Fisher Scientific, FD1133) prior to the electrophoresis, which converted the recombined S1P1 fragment to 180 bp.

### PDAC patient samples and exercise intervention

We enrolled 151 patients with potentially resectable pancreatic cancer into a randomized study of a prehabilitation home exercise intervention compared to usual care. The details of the trial have been published^26^. Briefly, patients in the exercise program arm were recommended to engaged in at least 150 minutes of moderate-to-vigorous intensity aerobic exercise per week (self-monitored using Fitbit activity trackers) and to perform at least 2 full-body strengthening exercise sessions per week (patients were issued a set of resistance bands and video and paper guides). Patients in the usual care group were also issued Fitbits to monitor their physical activity and received handouts with a full-body stretching guide and information about the benefits of being physically active before surgery. Patients were enrolled in this prehabilitation exercise program while receiving neoadjuvant chemotherapy and/or chemoradiation and completed the intervention period at the time of preoperative restaging visits. Of note, 79 of the 151 patients (52%) completed neoadjuvant treatment and underwent pancreatectomy. There were no significant differences in self-reported exercise minutes or Fitbit-measured physical activity between study arms. Thus, for the study of tumor vasculature reported here, we combined all patients from both arms for whom there were both surgical specimens and Fitbit data available (n=40). We calculated the median number of minutes per week of “fairly active” (moderate), “very active” (vigorous) and “active” minutes (the sum of “fairly active” and “very active” minutes which represents moderate-to-vigorous physical activity) as recorded by Fitbit, then split patients into two groups: below, and above or equal the median number of active minutes (130 minutes for active, and 66 minutes for fairly active and very active). The analyses of the surgical tumor specimen for the 40 study patients are described above.

### Statistical analysis

Statistical analysis was performed using SPSS software (IBM). Values in graphs are reported as mean ±SEM, median with box and whiskers or Log_2_ fold change. Shapiro-Wilk’s normality test was performed on datasets prior to statistical analysis. Nonparametric data were analyzed using an unpaired Mann– Whitney t test and parametric data were analyzed using unpaired Student t test. For analysis including more than 2 groups, the statistical comparisons were based on biological significance: Sed *vs* Ex 8 or Sed *vs* Ex 16; Sed *vs* Ex or Gem *vs* Ex+Gem.

## Results

### Aerobic exercise improves PDAC blood and lymphatic vasculature without impacting tumor mass and metastasis

To compare the impact of different aerobic exercise intensities on the vasculature of PDAC tumors in mice, we exercised KPC-4662 tumor-bearing mice on treadmills at either 8 m/min or 16 m/min (∼60% or ∼80% VO_2_max, respectively^27^; **Fig. 1a**). Neither exercise intensity changed tumor weight or peritoneal carcinomatosis as compared to sedentary mice (**Fig. 1b, c**). However, exercise at 16 m/min, but not at 8 m/min, increased the percentage of vessels with open lumens (23.2 vs 32.3%, p= 0.03) and elongated vessels (2.3 vs 4.5%; p=0.017), without changing the total vessel number or vessel density (**Fig. 1d-h**), compared to sedentary mice. Because exercise at 16 m/min effectively remodeled PDAC vasculature, we used this intensity for further analysis. Tumors from mice that exercised at 16 m/min had more perfused vessels than tumors from sedentary mice (p=0.033) (**Fig. 1l**) and significantly decreased vascular leakage (**Fig. 1m**). Tumor hypoxia and the coverage of vessels by pericytes were not changed by exercise (**Supplementary** Fig. 2a-f). Similar changes in vasculature were found in a second model of orthotopic PDAC, Hy15549. In Hy 15549-bearing mice, exercise at 16 m/min did not affect the number of total vessels, vessel density or elongated vessels (**Supplementary** Figure 3a-d), while it increased the percentage of open lumens and decreased vascular leakage compared to tumors of sedentary mice (**Supplementary** Figure 3a**, e-g**). Exercise did not change pericyte vessel coverage in Hy 15549 mice (**Supplementary** Fig. 3h, i).

**Figure 1.**
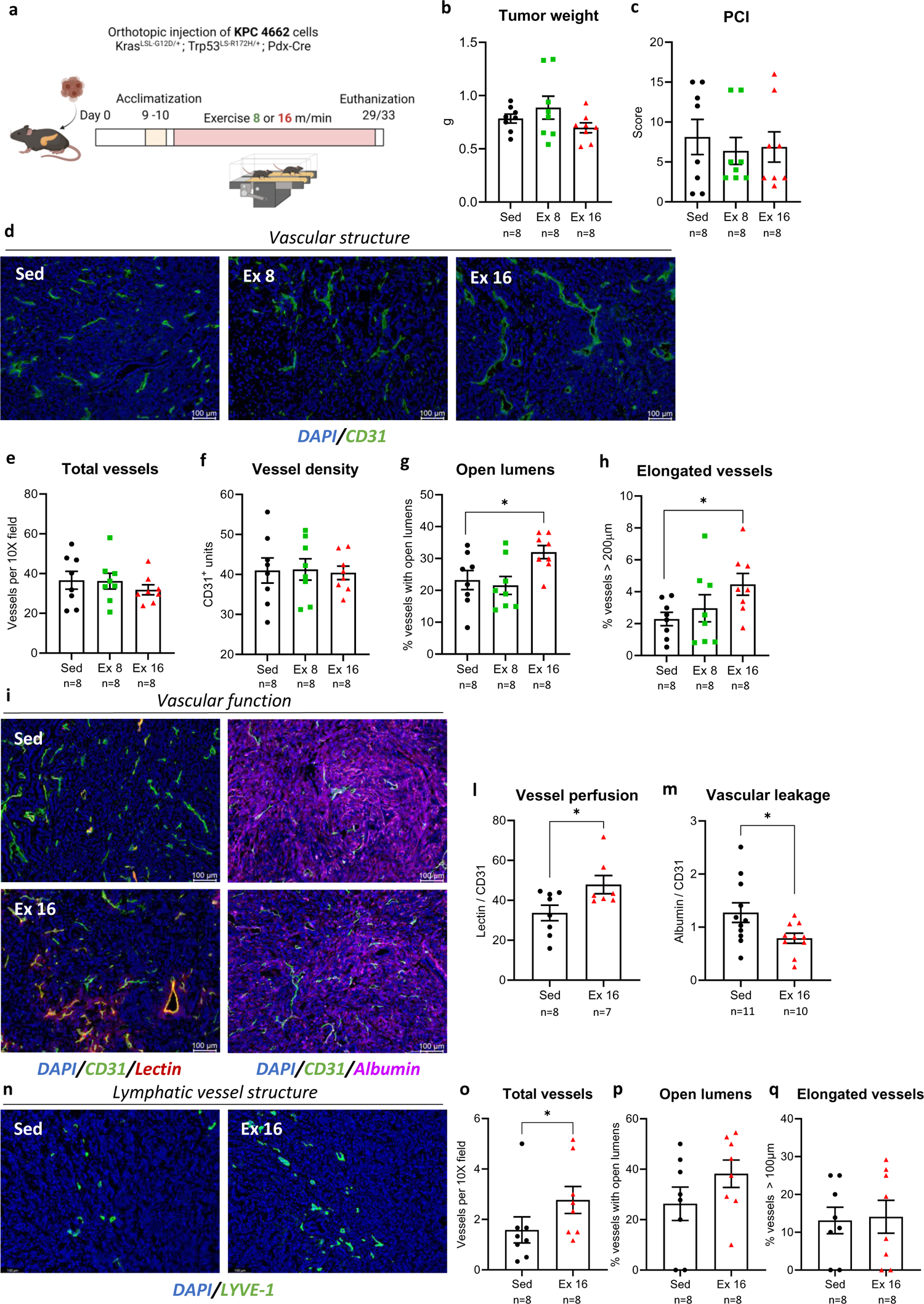
Aerobic exercise alone does not impact tumor mass or metastasis, but it improves tumor blood and lymphatic vasculature at higher intensity. a Experimental scheme of aerobic exercise training, including orthotopic injection of KPC-4662 cells, exercise acclimatization, exercise phase, and euthanization. b Tumor weights in grams (g). c Peritoneal carcinomatosis index (PCI) expressed as total metastasis score for each animal. d-h Representative images of tumor vascular structure. Vasculature was evaluated using immunofluorescence staining for CD31 (green) with DAPI (blue) counterstaining and quantified on five to nine sections per tumor. I-m Representative images of vessel perfusion and vascular leakage. Vessel perfusion was calculated as the ratio between lectin (red):CD31 (green) immunofluorescence signal. Vascular leakage was calculated as the ratio between albumin (magenta):CD31 (green) immunofluorescence signal. n-q Representative images of tumor lymphatic vessel structure. Lymphatic vessel structure was evaluated using LYVE-1 (green) immunofluorescence staining with DAPI (blue) counterstaining. Scale bar, 100μm. For all graphs, each point indicates one tumor. Bars indicate mean ± SEM. * p<0.05 relative to Sed.

Because lymphatic vasculature contributes to fluid homeostasis and interstitial fluid pressure which impacts drug delivery, lymphatic vessels were examined in the KPC-4662 model. Exercise significantly increased the number of lymphatic vessels (**Fig. 1n, o**) relative to tumors from control mice. Exercise did not change the percentage of lymphatic vessels with open lumens or elongated lymphatic vessels in tumors (**Fig. 1n, p, q**).

### Aerobic exercise increases chemotherapy delivery and efficacy in mouse PDAC

Blood and lymphatic vascular remodeling induced by exercise may increase the penetration of drugs within the tumor and thus improve chemotherapy efficacy. To test this, we treated KPC-4662-bearing mice with gemcitabine twice per week, with or without daily exercise at 16 m/min, for 3 weeks (**Fig. 2a**). Exercise alone or in combination with gemcitabine increased the percentage of elongated vessels and vessels with open lumens, but did not affect the total vessel number, vessel density or pericyte vessel coverage as compared to sedentary animals (**Fig. 2b-f**; **Supplementary** Fig. 4a, b). Exercise with or without gemcitabine also increased functional vasculature in tumors, as indicated by improved vessel perfusion and decreased vascular leakage, as compared to sedentary controls or mice treated only with gemcitabine (**Fig. 3a-c**). Consistent with improved vascular function leading to better delivery of chemotherapy, exercise increased gemcitabine concentration within the tumor (32% increase in exercised relative to gemcitabine alone treated mice; p= 0.025; **Fig. 3d**). The combination of exercise and gemcitabine was more effective in reducing both tumor growth (p=0.028) and peritoneal metastasis (62.5% reduction; p=0.002) as compared to gemcitabine alone (**Fig. 3e-g**).

**Figure 2.**
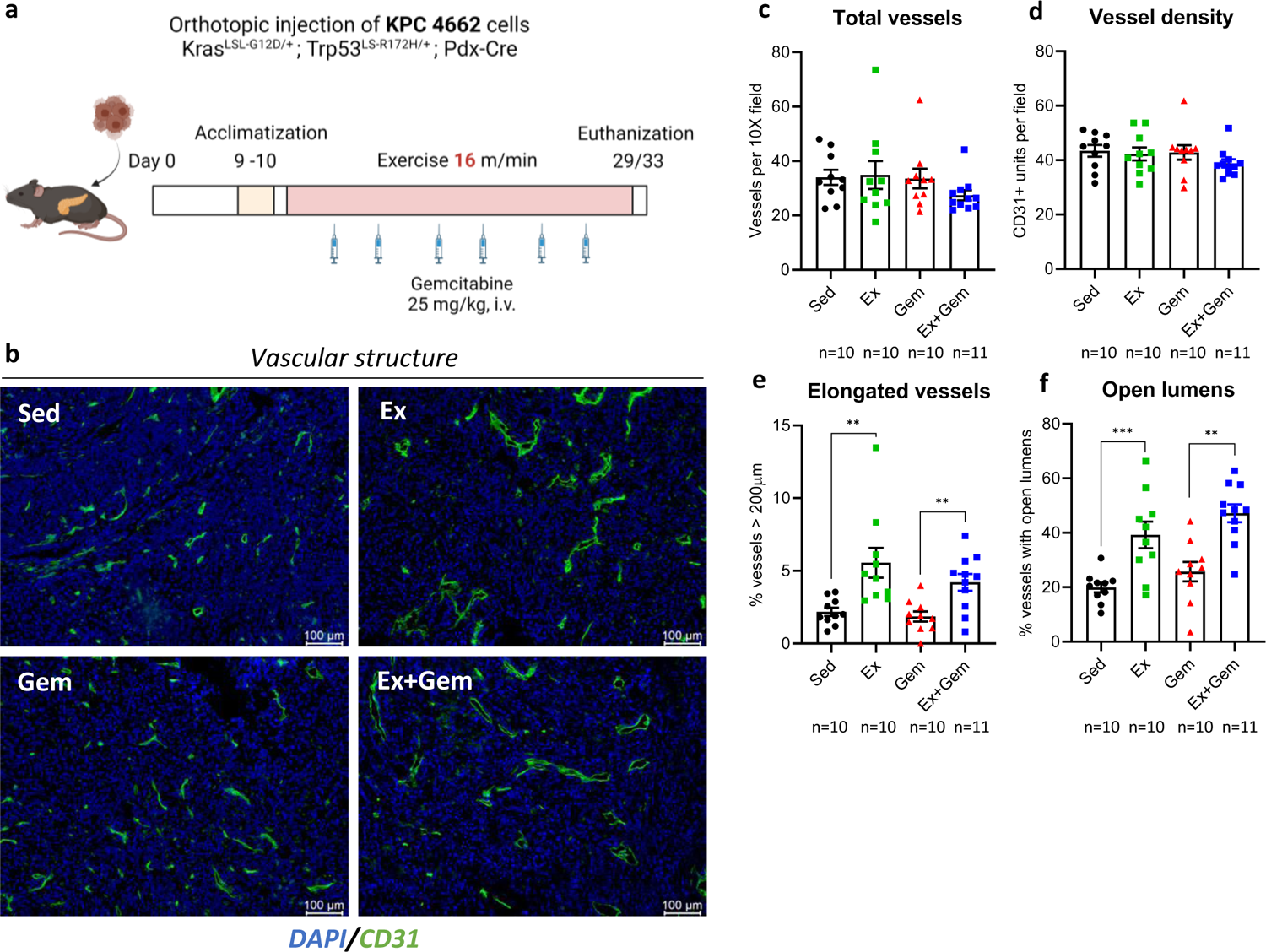
Exercise with or without chemotherapy improves tumor vascular structure. **a** Experimental scheme of aerobic exercise training, including KPC-4662 cells orthotopic injection, exercise acclimatization, gemcitabine treatment schedule and euthanization. **b-f** Representative images of tumor vascular structure. Tumor vascular structure was evaluated using CD31 (green) immunofluorescence staining with DAPI (blue) counterstaining. Scale bar, 100μm. For all graphs, each point indicates one tumor. Bars indicate mean ± SEM. ** p<0.01; *** p<0.001 relative to Sed or Gem.

**Figure 3.**
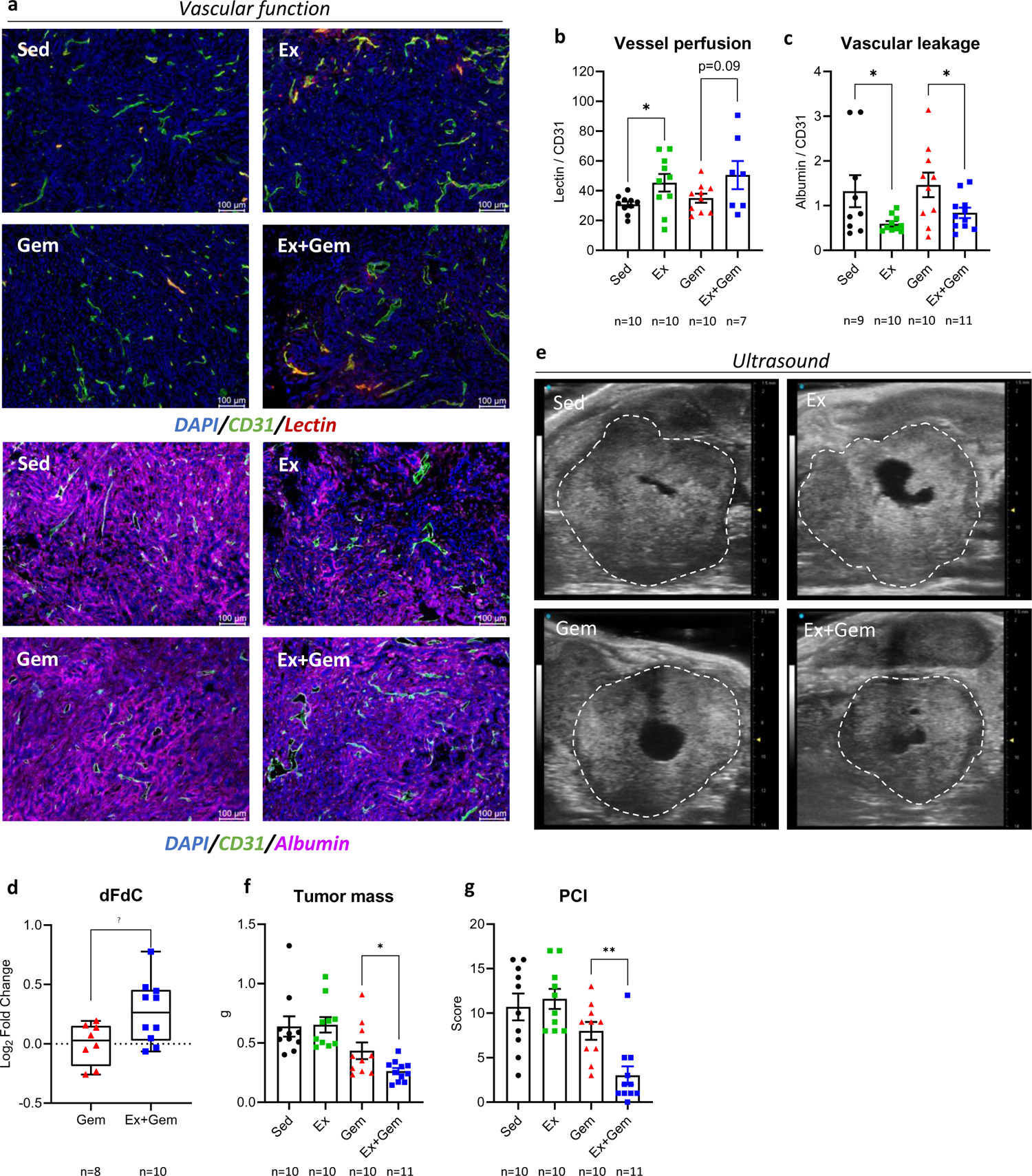
Exercise with or without gemcitabine improves vascular function, increases chemotherapy delivery and efficacy, and reduces peritoneal metastasis. **a-c** Representative images of vessel perfusion and vascular leakage. Vessel perfusion was calculated as the ratio between lectin (red):CD31 (green) immunofluorescence signal. Vascular leakage was calculated as the ratio between albumin (magenta):CD31 (green) immunofluorescence signal. Vascular leakage expressed as the ratio between albumin (purple):CD31 (green) immunofluorescence signal. Scale bar, 100μm. **d** Gemcitabine (dFdC) concentration measured by LC-MS and expressed as Log_2_ fold change *vs* Gem. **e, f** Representative ultrasound pictures of tumors showing difference in tumor area (dotted circle). Tumor weight in grams (g). **g** Peritoneal carcinomatosis index (PCI) expressed as total metastasis score for each animal. For all graphs, each point indicates one tumor. Bars indicate mean ± SEM, except for dFdC concentration expressed as median with box and whiskers. * p<0.05; ** p<0.01 relative to Sed or Gem.

### Aerobic exercise increases plasma S1P and activates S1PR1 on PDAC tumor endothelium

To determine whether improved tumor vascular function is necessary for the improved chemotherapy efficacy afforded by exercise, we first identified a key molecular mediator of the tumor vascular response to exercise, S1PR1. Exercise did not change the expression of S1PR1 on tumor endothelial cells (**Fig. 4a, b**). A reporter mouse model in which activation of S1PR1 is indicated by GFP expression (S1PR1-GFP mice^24^) was utilized to measure S1PR1 signaling in tumor endothelium. The fidelity of the GFP reporter in KPC-4662 tumors was confirmed by immunofluorescence (**Supplementary** Fig. 5a) and flow cytometry on CD31^+^ CD45^-^ endothelial cells (**Supplementary** Fig. 5b, c). Exercise caused a significant increase in S1PR1 activation in tumor endothelial cells compared to sedentary animals (**Fig. 4c**). Exercise also increased the plasma levels of the S1PR1 ligand, S1P, and its precursor sphingosine (**Fig. 4d, e**).

**Figure 4.**
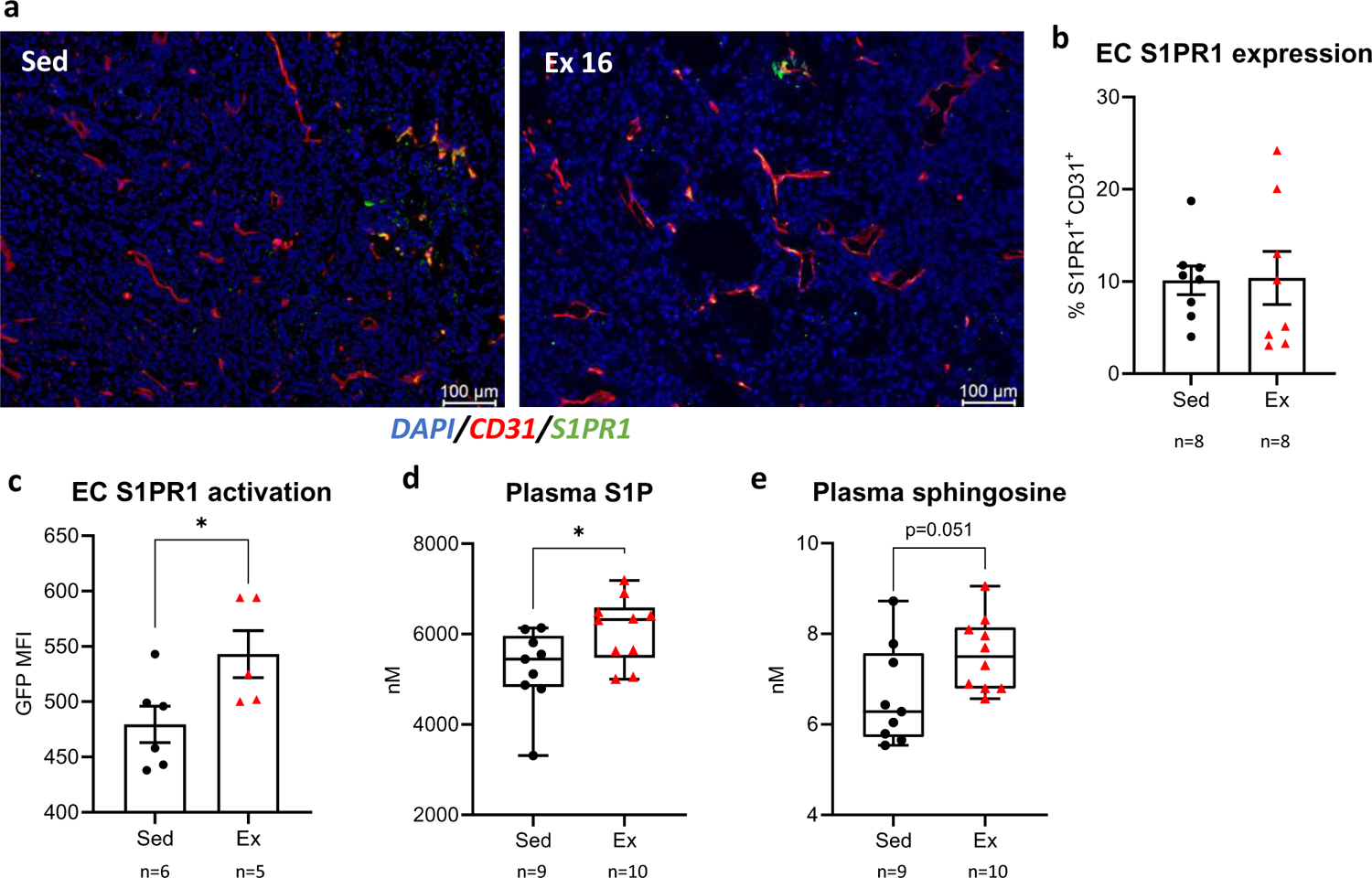
Aerobic exercise increases S1PR1 activity in tumor endothelial cells and plasma sphingosine and S1P, without affecting tumor endothelial cell S1PR1 expression. **a, b** Representative pictures of S1PR1 staining. S1PR1 tumor endothelial expression was analyzed by immunofluorescence and expressed as percentage colocalization between S1PR1 (red) and CD31 (green) immunofluorescence signal. Scale bar, 100μm. **c** Endothelial cell S1PR1 activation was analyzed by measuring GFP mean fluorescence intensity (MFI) by flow cytometry in CD31^+^ CD45^-^ cells. **d, e** Plasma sphingosine-1-phosphate (S1P) and sphingosine were analyzed by LC-MS and expressed as nM. For immunofluorescence and flow cytometry data graphs, each point indicates one tumor, bars indicate mean ± SEM. For LC-MS data, each point represents one mouse and summary data is shown as box and whiskers with median. * p<0.05 relative to Sed or Gem.

### S1PR1 expression is necessary for the exercise-dependent improvement in vascular function and chemotherapy efficacy

Exercise increased tumor endothelial cell S1PR1 activation, and improved tumor vascular function and chemotherapy efficacy. To define the role of S1PR1 in the exercise-induced remodeling of PDAC vasculature, we used a mouse model of conditional endothelial specific deletion of S1PR1^20^ (*S1pr1*^LoxP/LoxP^ *Cdh5*-*Cre*-ER^T2^; S1PR1 EKO). KPC-4662 tumors in S1PR1 EKO mice had increased tumor total endothelial cell area, vessel number and density, and decreased open lumens as compared to control animals (*S1pr1*^LoxP/LoxP^ without *Cdh5*-*Cre*-ER^T2^; S1PR1 WT; **Supplementary** Fig. 6a-e). In addition, S1PR1 endothelial KO increased vascular leakage, without changing vascular perfusion or pericyte vessel coverage in tumors (**Supplementary** Fig. 6f-i). Lymphatic endothelial cells also express *cdh5* and may be affected by the *cdh5-cre* driven *s1pr1* deletion^28^. Tumors from S1PR1 EKO mice had decreased total lymphatic vessels and percentage of elongated lymphatic vessels (**Supplementary** Fig. 6l-n). There was no difference in tumor mass and peritoneal metastasis between S1PR1 EKO and S1PR1 WT mice (**Supplementary** Fig. 6o, p).

Due to the activation of S1PR1 in tumor endothelium by exercise and the role of S1PR1 as an important mediator of vascular maturation and function, we hypothesized that S1PR1 may be necessary for improvements in tumor vascular function, and gemcitabine efficacy, by exercise. To test this hypothesis, tumor-bearing S1PR1 EKO mice were exercised and treated with gemcitabine in the same way as S1PR1 WT mice (**Fig. 5a**). In contrast to S1PR1 WT mice, exercise did not remodel tumor vasculature or improve tumor vascular function (**Fig. 5b-f**; **Fig 6 a-c**) and did not change pericyte coverage of vessels in S1PR1 EKO mice (**Supplementary** Fig. 4c, d). Exercise improved lymphatic vessel number and structure also in the absence of endothelial S1PR1 (**Supplementary** Fig. 6q-s). Along with the loss of vascular remodeling in S1PR1 EKO mice in response to exercise, the combination of exercise plus gemcitabine was ineffective in increasing chemotherapy concentration and did not reduce tumor growth and peritoneal metastasis as compared to gemcitabine alone in S1PR1 EKO mice (**Fig. 6d-e**). Taken together, these results suggest that improved vascular function mediated by S1PR1 is necessary for improved gemcitabine efficacy against PDAC in mice.

**Figure 5.**
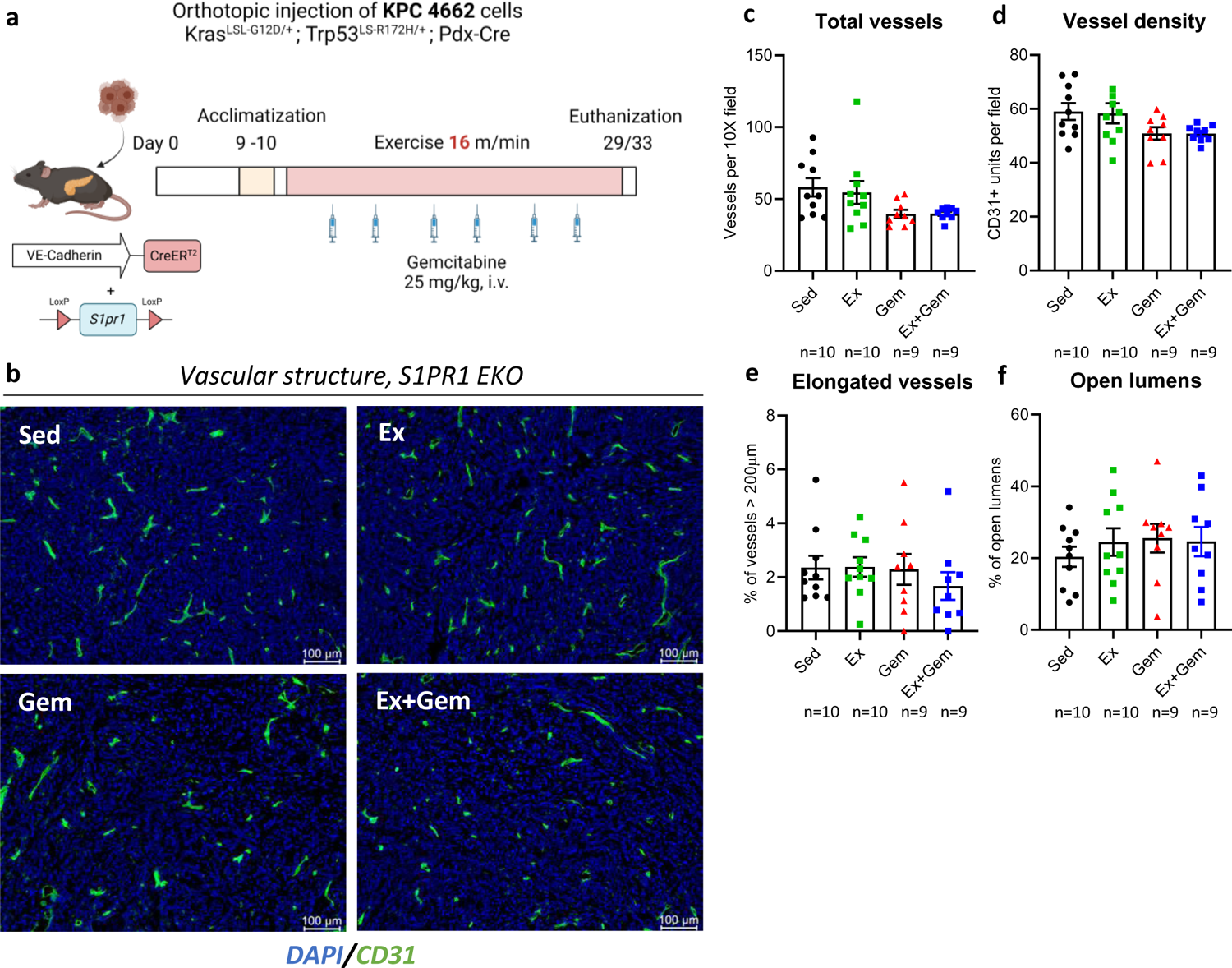
S1PR1 EKO blunts the exercise-induced improvements of tumor vascular structure. **a** Experimental scheme of aerobic exercise training, including KPC-4662 cells orthotopic injection, exercise acclimatization, gemcitabine treatment schedule and euthanization in S1PR1 EKO mice. **b-f** Representative pictures of S1PR1 staining. Tumor vascular structure was evaluated using CD31 (green) immunofluorescence staining with DAPI (blue) counterstaining. Scale bar, 100μm. For all graphs, each point indicates one tumor. Bars indicate mean ± SEM.

**Figure 6.**
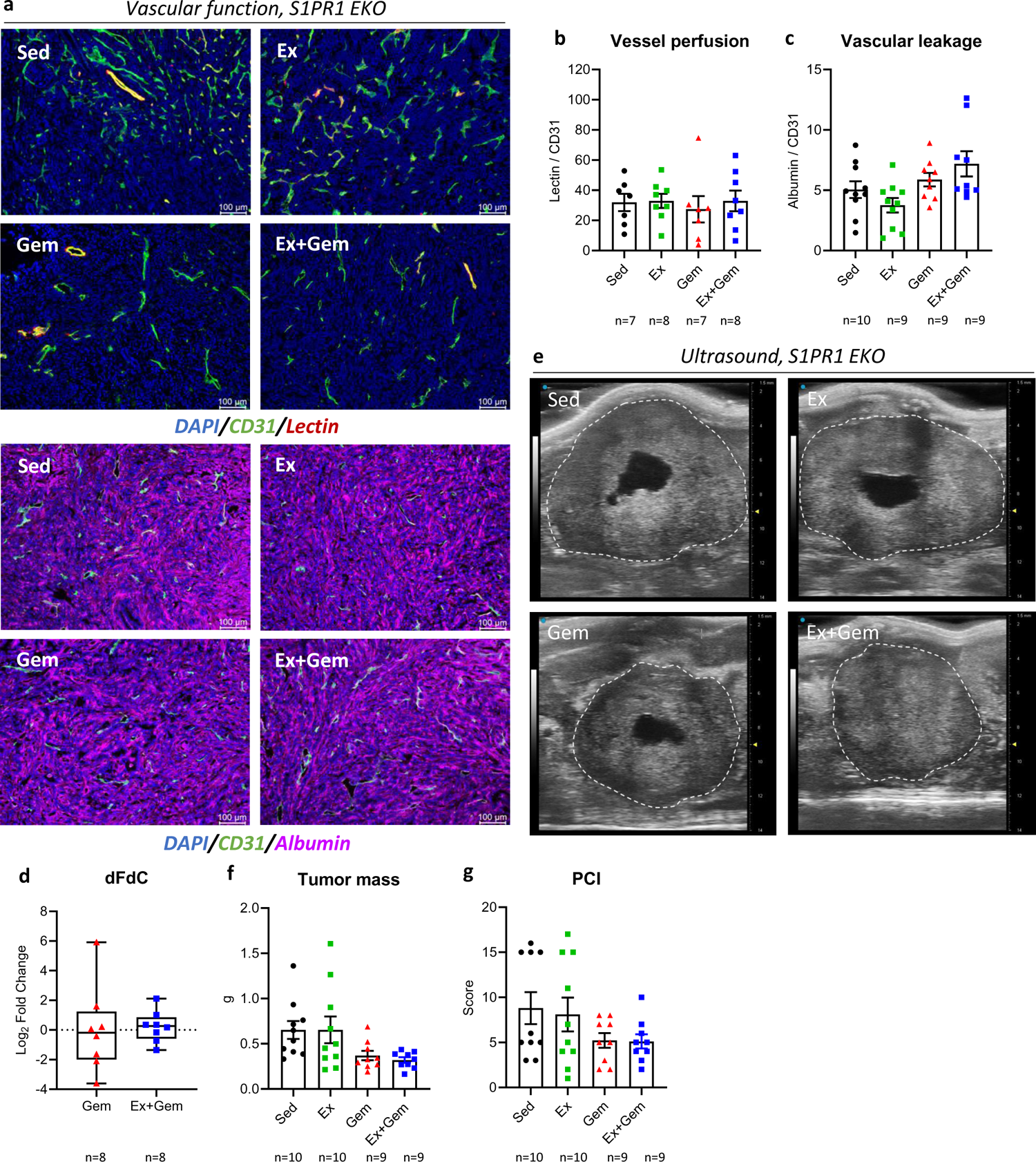
S1PR1 EKO blunts the exercise-induced improvements in tumor vascular function, chemotherapy delivery and efficacy and peritoneal metastasis. **a-c** Representative images of vessel perfusion and vascular leakage. Vessel perfusion was calculated as the ratio between lectin (red):CD31 (green) immunofluorescence signal. Vascular leakage was calculated as the ratio between albumin (magenta):CD31 (green) immunofluorescence signal. Scale bar, 100μm. **d** Gemcitabine (dFdC) concentration measured by LC-MS and expressed as Log_2_ fold change *vs* Gem. **e, f** Representative ultrasound pictures of tumors showing difference in tumor area (dotted circle). Tumor weight in grams (g). **g** Peritoneal carcinomatosis index (PCI) expressed as total metastasis score for each animal. For all graphs, each point indicates one tumor. Bars indicate mean ± SEM, except for dFdC concentration expressed as median with box and whiskers.

### Increased vigorous physical activity in PDAC patients remodels tumor vasculature and reduces tumor vascular leakage

We next validated the relevance of our findings in mice with PDAC tumors from patients who participated in a study of home-based exercise during neoadjuvant therapy. Study design, clinical demographics of participants, and primary outcomes were previously reported^26^. Participants were randomized to enhanced usual care or to prescribed aerobic and resistance exercise. Tumor specimens were obtained at the time of surgical resection for those patients who underwent surgery after neoadjuvant therapy. Despite the exercise intervention, objectively measured weekly activity (by Fitbit) and self-reported weekly moderate-to-strenuous physical activity minutes were not significantly different between study arms. Due to of the lack of significant differences in physical activity between the two arms, we also pooled the participants and divided them into two groups based on weekly fairly active, active or very active minutes (moderate, moderate-to-vigorous and vigorous physical activity, respectively; <median *vs* ≥median of average minutes/week) as measured by Fitbit. Tumors of patients that performed more than the median number of minutes per week of vigorous physical activity showed increased total vessel, vessel density, number of vessels with open lumens and elongated vessels (**Fig. 7a-e**). In the same patients, there was a significant decrease in tumor vascular leakage as indicated by albumin. Grouping the patients by experimental study arms or as above or below the median of weekly fairly active and active minutes did not show differences between groups, except for the increased number of vessels with open lumens in patients above the median number of active minutes per week (**Supplementary** Fig. 7).

**Figure 7.**
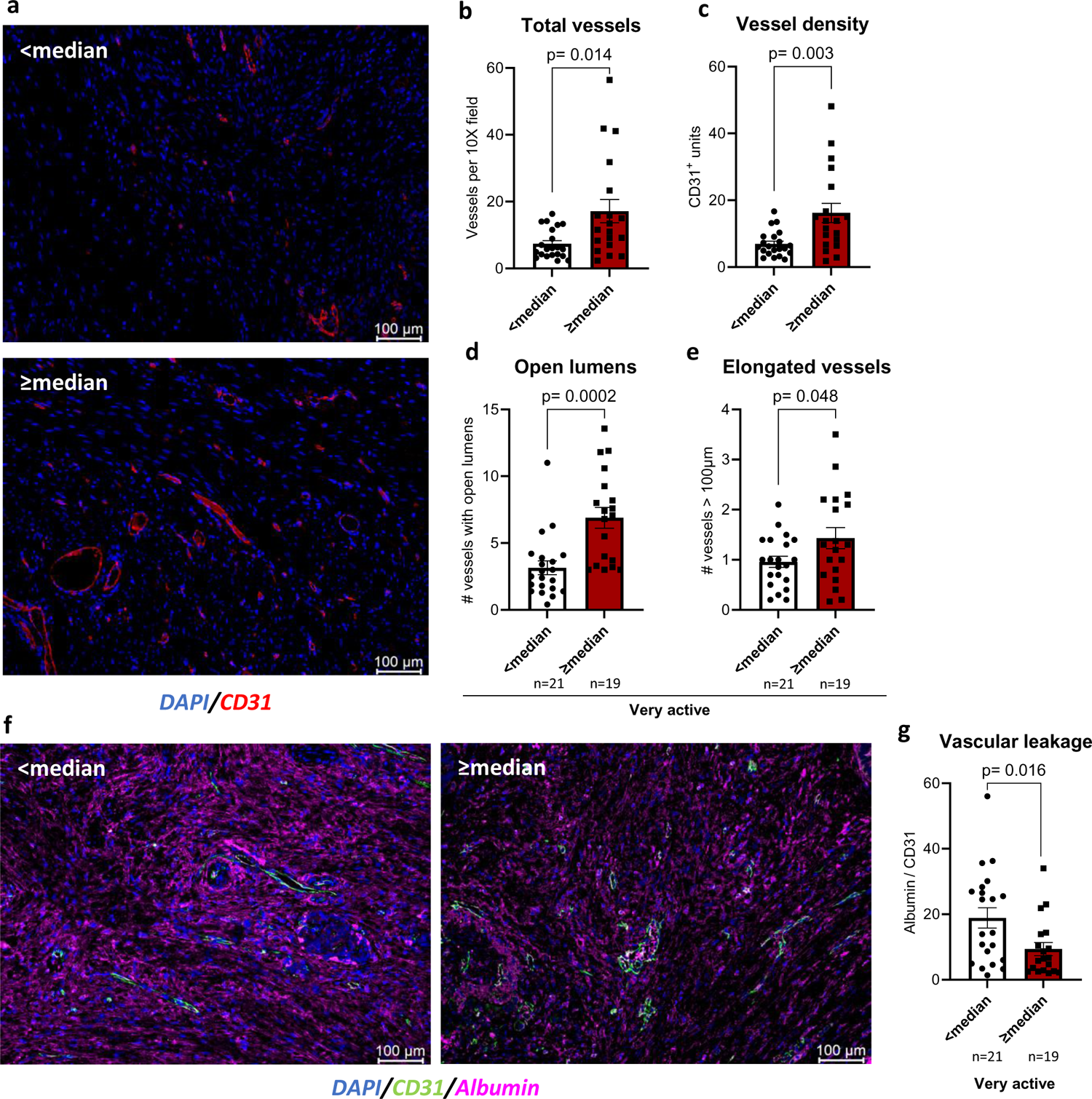
Exercise during neoadjuvant therapy remodeled tumor vasculature and decreased tumor vascular leakage in PDAC patients. Patients were divided in two groups based on the median number of minutes of very active physical activity performed weekly, as tracked by Fitbit (<66 *vs* ≥66 minutes). Tumors obtained during surgical resection were assessed by immunofluorescent staining. **a-e** Representative images of tumor vascular structure. Tumor vascular structure was evaluated using CD31 (red) immunofluorescence staining with DAPI (blue) counterstaining. **f, g** Representative images of vascular leakage. Vascular leakage was calculated as the ratio between albumin (magenta):CD31 (green) immunofluorescence signal. Scale bar, 100μm. Bars indicate mean ± SEM. p values are shown above bars.

## Discussion

Despite the advancements made in therapeutic approaches against PDAC, the majority of patients have disease that continues to progress on therapy, leading to low survival rates^29^. Chemotherapy efficacy against PDAC is blunted in part by poor delivery to the tumor cells. Drug delivery to the tumor is impeded by dysfunctional, sparse blood and lymphatic vasculature^30^, by the solid stress imposed by cancer cell growth, and by the excessive deposition of extracellular matrix that characterizes PDAC^31^. In this study, we demonstrated that aerobic exercise is an effective tool for improving chemotherapy delivery and efficacy against pancreatic tumors and peritoneal metastasis in mice. These effects were mediated by tumor vascular normalization via S1P – S1PR1 signaling. When blood vessels were unable to remodel in response to exercise, the added benefit of exercise for chemotherapy efficacy was lost. Evaluation of surgically resected PDAC from patients who participated in an exercise intervention during neoadjuvant therapy indicates that exercise may be a useful clinical tool to support microenvironment changes that allow better drug access to the tumor.

We did not find an effect of exercise without chemotherapy on tumor growth or metastasis. This is in contrast to other preclinical studies which found that exercise alone reduced tumor growth^13,32–35^. The effect of exercise on tumor growth varies by the tumor type, tumor location, exercise protocol, and the time of exercise initiation relative to tumor inoculation. Despite no effect of exercise alone on tumor growth, exercise combined with gemcitabine reduced the primary tumor growth and peritoneal metastasis more than gemcitabine alone. Exercise decreased tumor vessel permeability and increased gemcitabine concentration within tumors, consistent with reduced metastatic spread related to better drug delivery and/or an improved physical barrier to tumor cell extravasation. Exercise also increased the number of lymphatic vessels in the tumor. Although increased lymphatic vessels may have the potential to increase the dissemination of tumor cells^36–39^, there was no difference in peritoneal spreading of PDAC in exercised mice in the absence of chemotherapy. When delivered with gemcitabine, exercise reduced peritoneal metastasis of PDAC in mice. The reduction of PDAC metastasis is a novel finding of critical importance because when patients develop metastatic disease during neoadjuvant therapy, they become ineligible for surgery. Thus, our work supports a potential role for exercise during neoadjuvant therapy to increase the chance of curative surgery.

Aerobic exercise has been shown to improve anti-cancer therapy in different animal models^11,12,14,33,40–42^. While some studies have suggested that this is due to improved blood vessel function and one has demonstrated a role for Thrombospondin-1 in the tumor endothelial response to exercise^11^, the specific regulation of tumor vasculature by exercise and the relevance of these changes remain enigmatic. We found that exercise increased S1PR1 activation on tumor endothelium, consistent with the known function of S1PR1 in promoting improved tumor vascular structure and function^23^. Endothelial cell-specific *s1pr1* deletion prevented the exercise induced improvements of tumor blood vessels but not lymphatic vessels, indicating that exercise-induced blood, but not lymphatic, vessel remodeling is dependent on S1PR1. Exercise did not increase gemcitabine concentration and efficacy in S1PR1 EKO mice, indicating a causal relationship of improved tumor blood vessel function leading to increased chemotherapy delivery and efficacy. Because the effects of exercise in lymphatic vessels were not different in S1PR1 EKO relative to S1PR1 WT mice, we conclude that an improvement in blood vessels alone is sufficient for exercise-induced increased chemotherapy delivery and efficacy. In summary, the present study defined S1PR1 as a necessary mediator of the PDAC vascular remodeling and increased chemotherapy efficacy upon exercise.

Exercise is a feasible and effective tool to improve physical fitness and quality of life in patients undergoing treatment for PDAC^26,43,44^. Additionally, improved tumor vessel maturity and function correlates with better outcomes for patients with PDAC who receive neoadjuvant chemotherapy^45^. Here, we demonstrated in mice that exercise improved blood vessel function, improved chemotherapy efficacy, and decreased metastasis. We also validated the relationship between vigorous physical activity and tumor vasculature in patient PDAC using a cohort from a clinical study in which patients underwent an exercise program during neoadjuvant therapy. The limitations of this clinical study include the inaccuracy of the activity tracker to measure the prescribed exercise program rather than day-to-day activity, and the lack of difference in self-reported exercise minutes or Fitbit-measured physical activity between study arms, leading us to divide the patients into groups based on the median of weekly Fitbit-measured activity minutes. Taken together, our data in mice and in patients suggests that exercise may be beneficial for increasing the efficacy of anti-cancer treatment and survival.

## Acknowledgments

R.B., J.L. and K.S. were supported by Cancer Prevention and Research Institute of Texas (CPRIT, grant RP190256). The clinical trial was supported by a generous gift from the Wyck Knox Family Foundation. P.T. was supported by CPRIT, grant RP210028. S.P. was supported by the Pauline Altman-Goldstein Foundation Discovery Fellowship.

## Author Contributions

R.B. contributed to conceptualization, design, data acquisition, data curation, formal analysis, validation, investigation, methodology, writing–original draft, writing–review and editing. S.P., J.L. and H.P. contributed to data acquisition, formal analysis, data curation, writing–review and editing. P.T. and K.H. contributed to formal analysis, data curation, writing–review and editing. G.L. contributed to formal analysis and data curation. H.S. and B.G. contributed to conceptualization, writing–review and editing. A.N., N.P., M.H.K., M.P. and F.M. conducted the clinical trial and provided access to patient samples. R.W. contributed to data curation, writing–review and editing. T.L. and P.L.L. contributed to the analysis and writing–review and editing. K.S. contributed to conceptualization, resources, data curation, formal analysis, supervision, funding acquisition, writing–original draft, project administration, writing–review and editing.

## Materials & correspondence

Keri Schadler, PhD. The University of Texas MD Anderson Cancer Center, 1515 Holcombe Blvd, Houston, TX 77030. Phone: 713-794-1035; Fax: 713-563-5604;

---

**Supplementary Figure 1. S1PR1 EKO genotype verification, tumor volume calculation and experimental scheme. a** Lung DNA characterization from *S1pr1*^flox/flox^, *S1pr1*^flox/flox^ Cdh5-Cre-ER^T2^ (S1PR1 EKO) and wild-type (WT) mice. The combination of primers used amplified different products: 250 bp S1PR1 loxP; 200 bp S1PR1 WT or KO; 180 bp SacI digested S1PR1 KO fragment ΔEx2. All mice were treated with tamoxifen as described in methods, except for WT. **b** Representative images of S1PR1 (green) immunofluorescence staining in CD31^+^ (red) vessels in S1PR1 WT (*S1pr1*^flox/flox^) and S1PR1 EKO mice. Scale bar, 50 μm. **c** Representative tumor ultrasound pictures. For tumor volume calculation, two different pictures of a single tumor were acquired by ultrasound. The pictures were obtained taking two tangential ultrasound images along the transverse and longitudinal planes. Tumor volume was calculated by the formula 4/3πABC, where A, B and C are the measured diameters of each tumor. Values were expressed as mm^3^. **d** Exercise, acclimatization and ultrasound (US) experimental scheme for 8 and 16 meters/minute (m/min) exercise protocols.

**Supplementary Figure 2. Exercise does not impact hypoxia or pericyte vessel coverage in tumors**. **a, b** Representative images of tumor hypoxia. Tumor hypoxia was calculated as the ratio between hypoxyprobe (green) immunofluorescence signal and the number of DAPI^+^ (blue) nuclei. Scale bar, 1 mm. **c-f** Representative images of perycite vessel coverage. Pericyte vessel coverage was evaluated by analyzing the percentage of colocalization between αSMA^+^, desmin^+^ or NG2^+^ pericyte (magenta) and CD31^+^ (green) vessels, with DAPI (blue) counterstaining. Scale bar, 100μm. For all graphs, each point indicates one tumor. Bars indicate mean ± SEM.

**Supplementary Figure 3. Exercise at 16 m/min remodels vascular structure and decreases vascular leakage in Hy 15549 tumors. a-e** Representative images of tumor vascular structure. Tumor vascular structure was evaluated using CD31 (green) immunofluorescence staining with DAPI (blue) counterstaining. **f, g** Representative images of vascular leakage. Vascular leakage was calculated as the ratio between albumin (magenta):CD31 (green) immunofluorescence signal. **h, i** Pericyte vessel coverage was evaluated by analyzing the percentage of colocalization between markers αSMA^+^ or NG2^+^ pericyte (magenta) and CD31^+^ (green) vessels, with DAPI (blue) counterstaining. Scale bar, 100μm. For all graphs, each point indicates one tumor. Bars indicate mean ± SEM. * p<0.05 relative to Sed.

**Supplementary Figure 4. Exercise with or without chemotherapy does not affect pericyte vessel coverage in S1PR1 WT or EKO mice. a-d** Pericyte vessel coverage was evaluated by analyzing the percentage of colocalization between αSMA^+^ or NG2^+^ pericyte (magenta) and CD31^+^ (green) vessels, with DAPI (blue) counterstaining. Scale bar, 100μm. Data are expressed as mean ± SEM.

**Supplementary Figure 5. S1PR1-GFP reporter mouse verification and flow cytometry gating strategy. a** Representative image of KPC-4662 tumors from H2B-GFP (control) or S1PR1-GFP mice; DAPI^+^ (blue) GFP^+^ (green) signal in CD31^+^ (red) vessels. White arrows indicate GFP-positive endothelial cell nuclei. Scale bar, 100μm. **b** Gating strategy for the flow cytometry analysis of S1PR1 activation in tumor endothelial cells of S1PR1-GFP reporter mice. Single cell suspensions are gated for CD31^+^ and CD45^-^ cells for the identification of vascular endothelial cells. **c** Representative histogram showing the GFP fluorescence intensity for H2B-GFP (red) and S1PR1-GFP (green) tumors.

**Supplementary Figure 6. S1PR1 EKO alters blood and lymphatic vessel structure and vascular leakage. a** Tumor CD31 immunofluorescence area was expressed as arbitrary units and evaluated in tile scan tumor sections. **b-e** Tumor vascular structure was analyzed using CD31 immunofluorescence staining with DAPI counterstaining. **f** Vessel perfusion was calculated as the ratio between lectin (red):CD31 (green) immunofluorescence signal. **g** Vascular leakage was calculated as the ratio between albumin (magenta):CD31 (green) immunofluorescence signal. **h-i** Pericyte vessel coverage was evaluated by analyzing the percentage of colocalization between αSMA^+^ or NG2^+^ pericyte and CD31^+^ vessels, with DAPI counterstaining. **l-n, q-s** Lymphatic vessel structure was evaluated using LYVE-1 immunofluorescence staining with DAPI counterstaining in S1PR1 WT *vs* EKO mice or S1PR1 KO Sed *vs* Ex. **o** Tumor weight was expressed as grams (g). **p** Peritoneal carcinomatosis index (PCI) was expressed as total metastasis score for each animal. For all graphs, each point indicates one tumor. Bars indicate mean ± SEM. * p<0.05; ** p<0.01 relative to S1PR1 WT.

**Supplementary Figure 7.** There are no significant changes in tumor vascular structure and tumor vascular leakage when patient samples are divided by study arms, or by the median of weekly fairly active or active minutes. a-n Tumor vascular structure was evaluated using CD31 (red) immunofluorescence staining with DAPI (blue) counterstaining. o, q Vascular leakage was calculated as the ratio between albumin (magenta):CD31 (green) immunofluorescence signal. Scale bar, 100μm. Patients were divided in two groups based on arms (Arm A = Usual Care and Arm B = Exercise; or based on the median number of minutes of fairly active or active physical activity performed weekly (<66 *vs* ≥66 minutes and <130 *vs* ≥130 minutes, respectively). Bars indicate mean ± SEM. p values are shown above bars.

