## Supplementary Figures for "Aerobic exercise promotes PDAC vascular normalization through S1PR1 signaling in tumor endothelial cells"

### Supplementary Figure 1

a

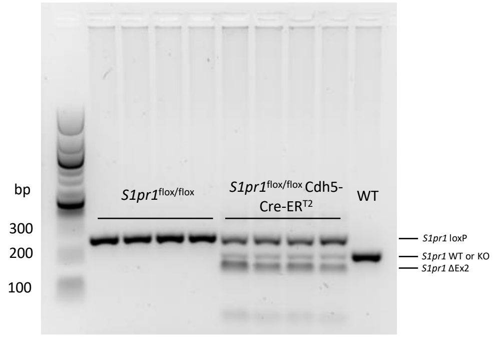


b

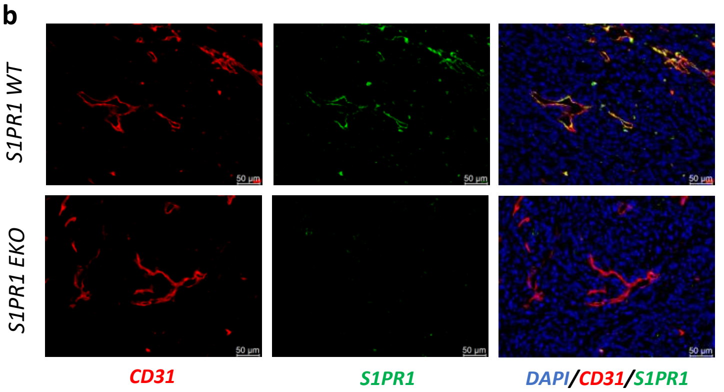


c

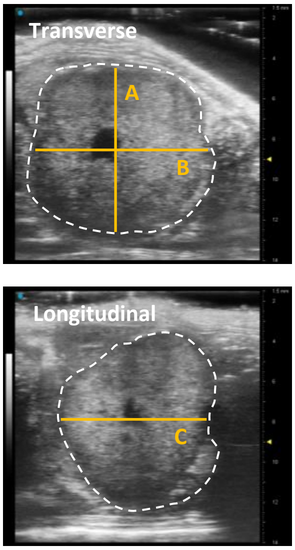


d

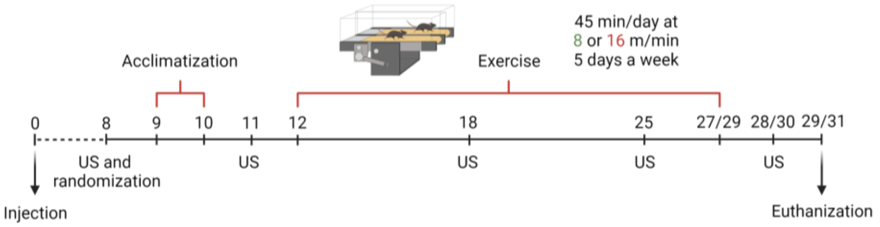


| Acclimatization Schedule | Day | 9 | 10 | 11 | 12 |
|---|---|---|---|---|---|
| 8 m/min | Intensity (m/min) | 3 | 6 | / | 8 |
| | Time | 20 | 30 | / | 45 |
| 16 m/min | Intensity (m/min) | 8 | 12 | / | 16 |
| | Time | 20 | 30 | / | 45 |

### Supplementary Figure 2

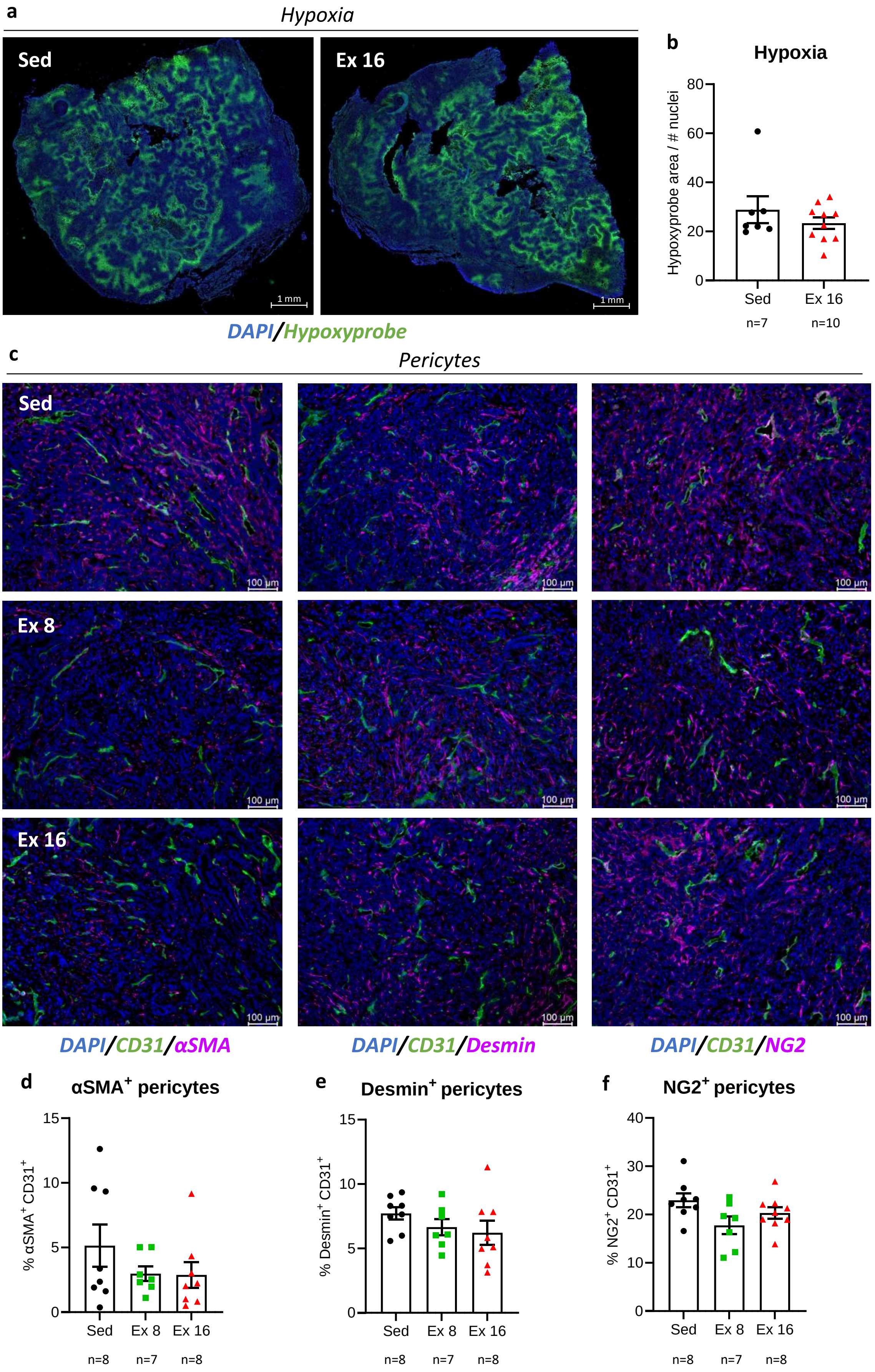

### Supplementary Figure 3

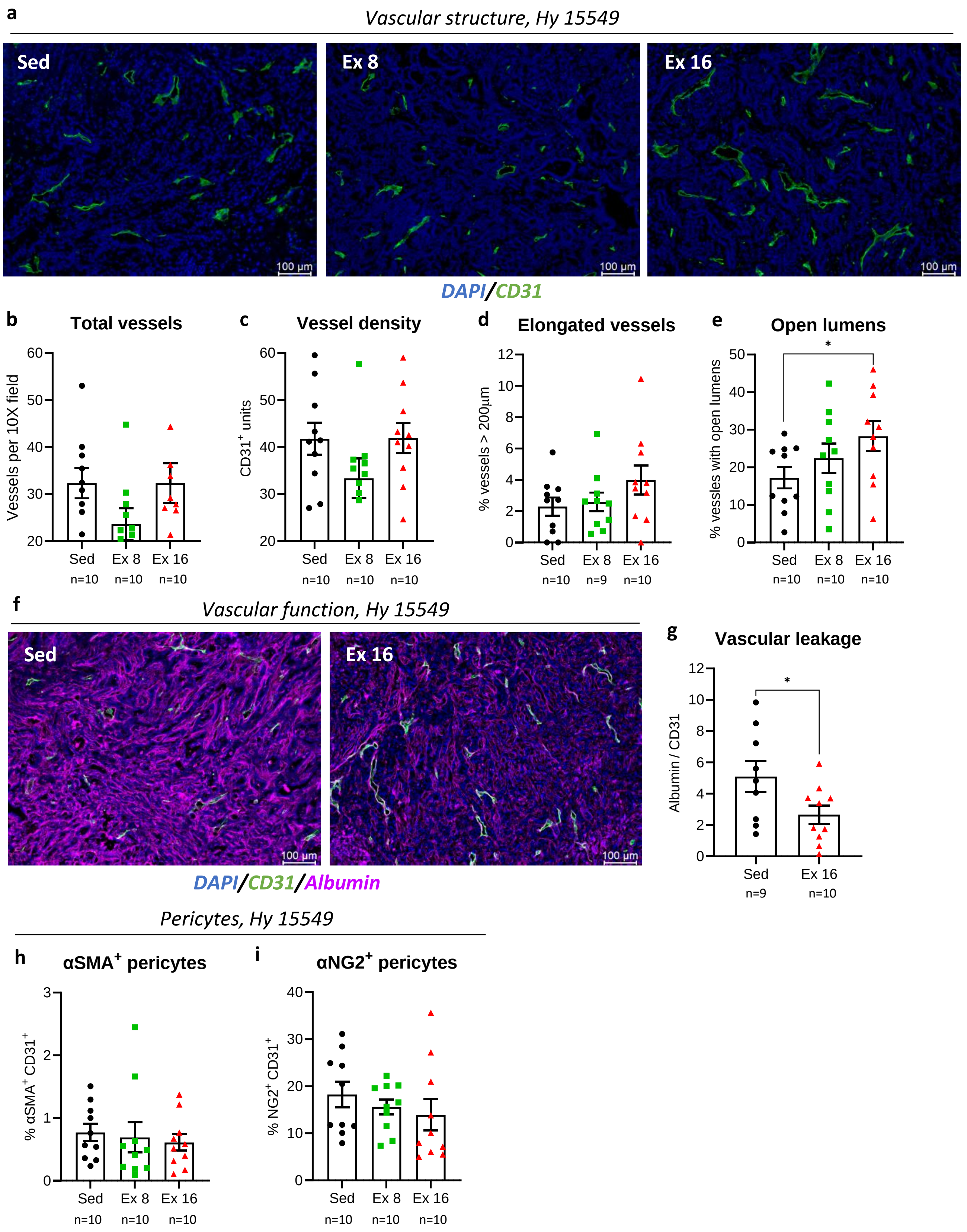

### Supplementary Figure 4

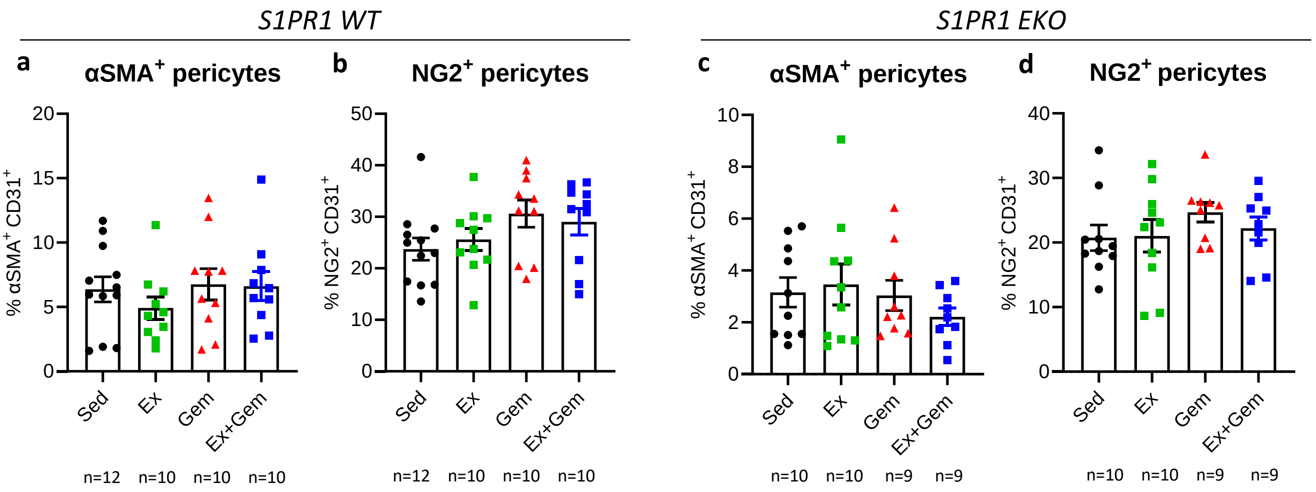

**Supplementary Figure 5**

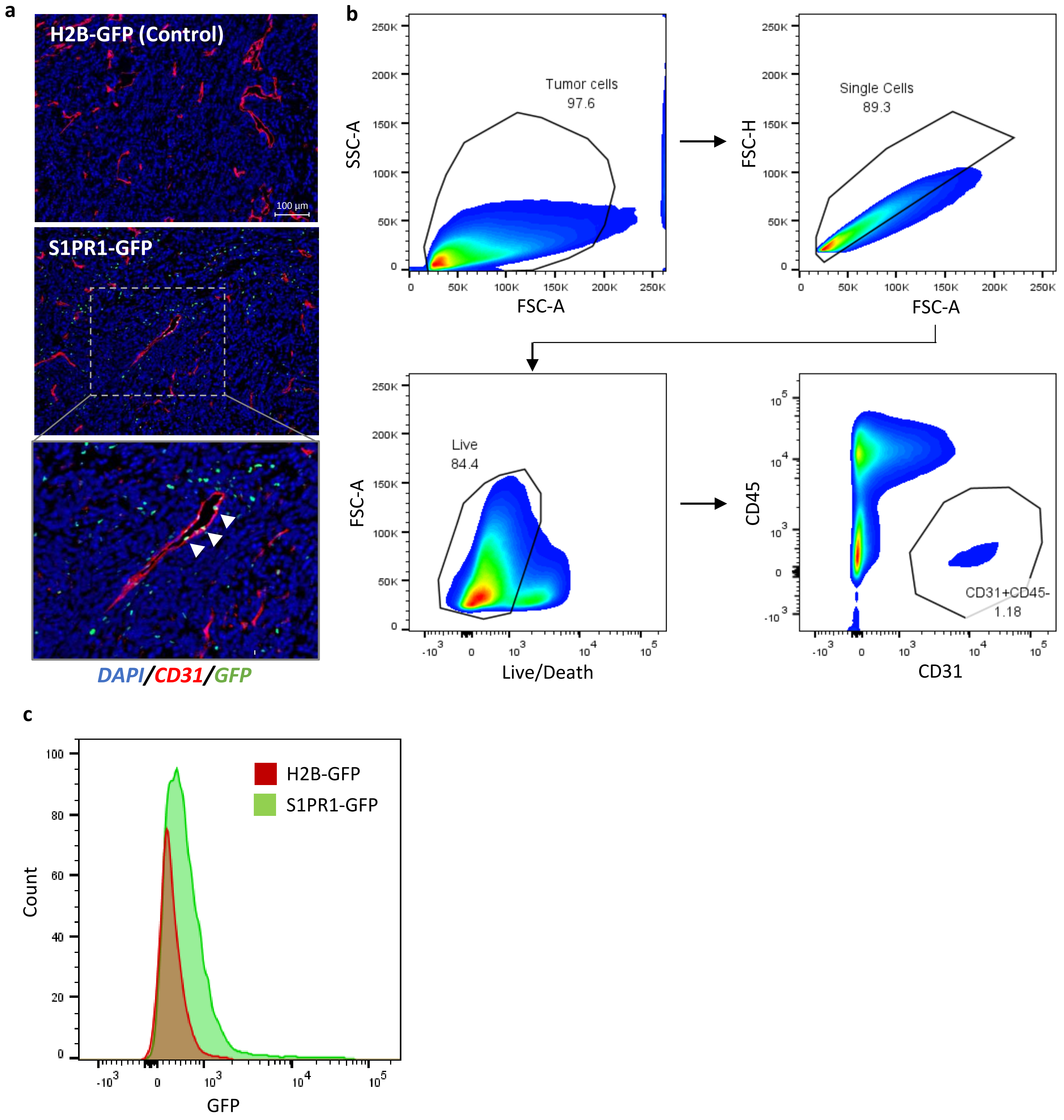

### Supplementary Figure 6

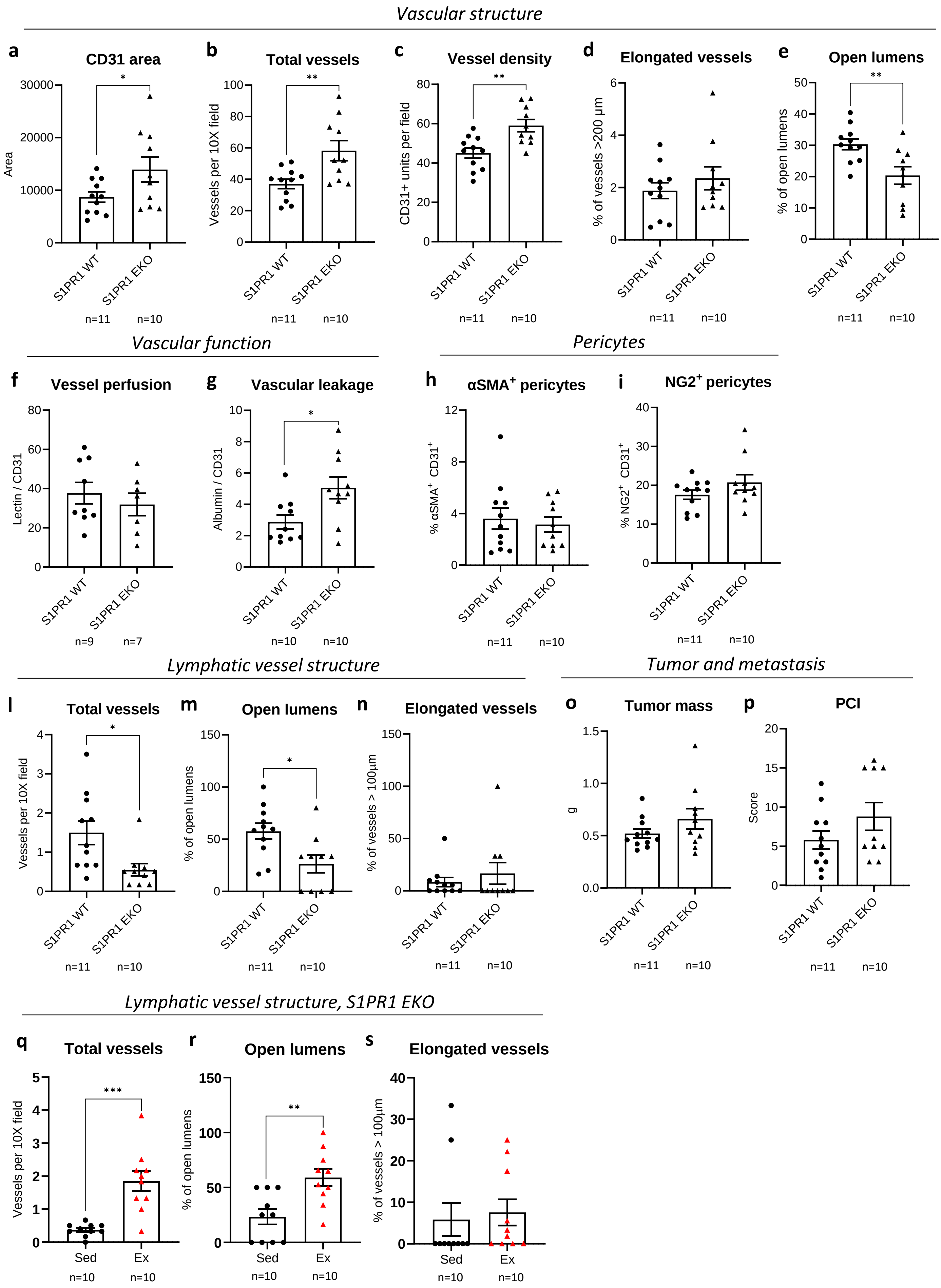

### Supplementary Figure 7

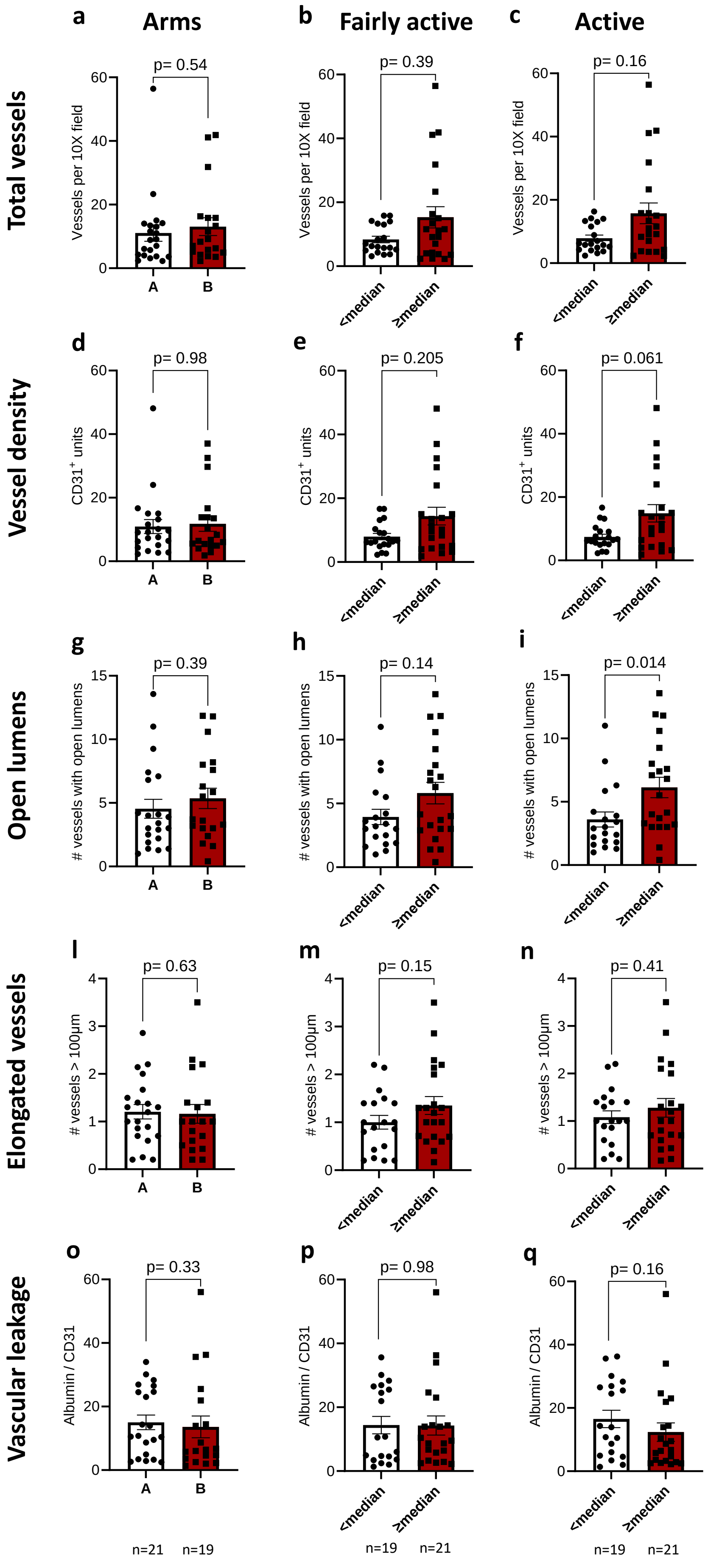
